# Static length changes of cochlear outer hair cells can tune low-frequency hearing

**DOI:** 10.1101/228353

**Authors:** Nikola Ciganović, Rebecca L. Warren, Batu Keçeli, Stefan Jacob, Anders Fridberger, Tobias Reichenbach

## Abstract

The cochlea not only transduces sound-induced vibration into neural spikes, it also amplifies weak sound to boost its detection. Actuators of this active process are sensory outer hair cells in the organ of Corti, whereas the inner hair cells transduce the resulting motion into electric signals that propagate via the auditory nerve to the brain. However, how the outer hair cells modulate the stimulus to the inner hair cells remains unclear. Here, we combine theoretical modeling and experimental measurements near the cochlear apex to study the way in which length changes of the outer hair cells deform the organ of Corti. We develop a geometry-based kinematic model of the apical organ of Corti that reproduces salient, yet counter-intuitive features of the organ’s motion. Our analysis further uncovers a mechanism by which a static length change of the outer hair cells can sensitively tune the signal transmitted to the sensory inner hair cells. When the outer hair cells are in an elongated state, stimulation of inner hair cells is largely inhibited, whereas outer hair cell contraction leads to a substantial enhancement of sound-evoked motion near the hair bundles. This novel mechanism for regulating the sensitivity of the hearing organ applies to the low frequencies that are most important for the perception of speech and music. We suggest that the proposed mechanism might underlie frequency discrimination at low auditory frequencies, as well as our ability to selectively attend auditory signals in noisy surroundings.

**Author summary:** Outer hair cells are highly specialized force producers inside the inner ear: they can change length when stimulated electrically. However, how exactly this electromotile effect contributes to the astonishing sensitivity and frequency selectivity of the inner ear has remained unclear. Here we show for the first time that static length changes of outer hair cells can sensitively regulate how much of a sound signal is passed on to the inner hair cells that forward the signal to the brain. Our analysis holds for the apical region of the inner ear that is responsible for detecting the low frequencies that matter most in speech and music. This shows a mechanisms for how frequency-selectivity can be achieved at low frequencies. It also opens a path for the efferent neural system to regulate hearing sensitivity.

## Introduction

Our ability to hear is due to an intricate mechanotransduction process that takes place inside the inner ear. Sound-evoked waves on the basilar membrane, an elastic structure stretching along the cochlear canal, cause the deflection of mechanosensitive hair bundles of the sensory cells, thus gating ion channels in the cell membrane and producing electrical signals that are ultimately transmitted to the brain [1]. The transfer of basilar-membrane motion to deflection of the hair bundles is shaped by the structurally complex organ of Corti (Fig. 1(A)), the outer hair cells of which can provide mechanical force [2]. Changes in transmembrane voltage cause these cells to change length, a phenomenon referred to as electromotility [3,4]. Furthermore, the hair bundles of outer hair cells can also generate mechanical force [5,6]. Both mechanisms may contribute to an active modulation of the sound-evoked motion of the organ of Corti [7–9].

**Fig 1.**
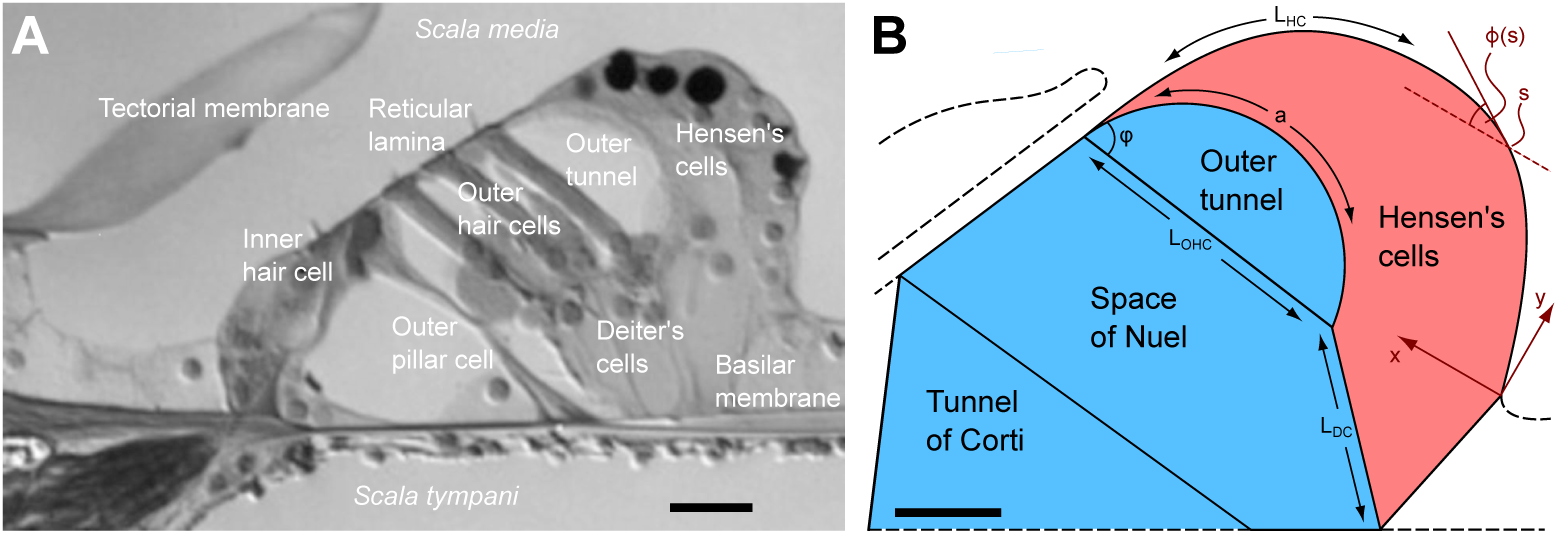
The organ of Corti and model geometry. (**A**) Micrograph of the apical organ of Corti from a guinea-pig cochlea [15]. Dark lipid droplets inside the Hensen cells serve as reflectors for a laser-interferometric beam. (**B**) Schematic representation of the organ of Corti as used in our geometric model. Length changes of the outer hair cell yield a deformation of the fluid space consisting of the tunnel of Corti, the space of Nuel, and the outer tunnel (blue) as well as the space of the body of Hensen cells (red) such that their cross-sectional areas are conserved separately. The scale bar denotes 20 *μ*m.

The active mechanical feedback by outer hair cells is essential for the extraordinary sensitivity, tuning, and dynamic range of mammalian hearing organs, and damage to the outer hair cells consequently results in hearing loss [10–12]. Although this feedback presumbaly operates on a cycle-by-cycle basis, outer hair cells can also exhibit quasi-static length changes that occur on much slower timescales. In particular, outer hair cells respond to acoustic stimulation through static contraction and sometimes elongation. Moreover, the static length change is largest when the frequency of the stimulation matches the characteristic frequency of the cochlear location of the outer hair cell [13,14]. Although discovered almost thirty years ago, the biophysical relevance of such static length changes of outer hair cells for the functioning of the organ of Corti remains unresolved.

Another main uncertainty regarding the feedback by outer hair cells concerns the micromechanics of the organ of Corti in the low-frequency apical region of the cochlea that is responsible for detecting frequencies below a few kHz that are most important for speech and music [16,17]. Recent *in vitro* experimental studies have indeed shown that the apical organ of Corti deforms in a complex and unexpected way [17–22]. When stimulated electrically, the outer hair cells contracted and pulled the reticular lamina, in which the hair bundles of outer hair cells are anchored, towards the basilar membrane. Surprisingly, the lateral portion of the organ of Corti composed of the Hensen cells moved in the opposite direction, away from the basilar membrane, at an amplitude larger than that of the reticular lamina [21]. No vibration could be detected from the adjacent portion of the basilar membrane [17]. The mechanisms producing this complex motion of the organ remain unclear.

Here we set out to identify the origin of the complex internal motion of the organ of Corti at the cochlear apex and the influence of static length changes of outer hair cells. We show that a plausible assumption about the apical organ of Corti, namely that each cross-section is incompressible, highly constrains the organ’s internal motion. The deformation of the organ of Corti that results from length changes of the outer hair cells can then be described through a mathematical model that is based on the organ’s geometry. We develop this model and verify it through comparison with existing [17,21] as well as newly acquired experimental data, where length changes of the outer hair cells were induced by current injections inside scala media. Our results reveal that static length changes of the outer hair cells can sensitively determine how much of the sound-evoked motion is transferred to the reticular lamina, thus providing a novel mechanism for outer hair cells to regulate hearing sensitivity.

## Results

### Local incompressibility of the apical organ of Corti

Sound elicits a traveling wave on the basilar membrane which triggers the deflection of hair bundles and thus the electromotile response of the outer hair cells. As the outer hair cells contract, the reticular lamina and basilar membrane are pulled towards each other [7] (Fig. 1). This can potentially reduce the cross-sectional area of the fluid-filled space of Nuel, causing fluid inside the organ of Corti to be displaced longitudinally, that is, along the cochlear canal. The volume of displaced fluid is proportional to the change in cross-sectional area, multiplied by the longitudinal extent *l* of the organ that contracts. For a traveling wave, this longitudinal extent *l* is approximately half the wavelength, and the amplitude of the evoked fluid velocity is proportional to the displaced volume and thus to the wavelength.

Near the cochlear apex, low-frequency sound elicits a wave with a long wavelength of several millimeters [2,23]. The longitudinal extent over which the organ of Corti deforms similarly thus far exceeds the width and the height of the space of Nuel, which are of the order of 100 *μ*m. Longitudinal fluid flow inside the organ of Corti would thus require velocities much larger than the velocity of the length-changing outer hair cells, and would hence be counteracted by viscous friction. We conclude that such longitudinal flow is suppressed and that the cross-sectional area of the organ of Corti in the apex is accordingly conserved when the outer hair cells change length. The same reasoning holds for *in vitro* experiments using electrical stimulation, due to the long effective range of the electrodes, but not for deformation of the organ of Corti near the cochlear base where the wavelengths in the peak region of a traveling wave can be much shorter, below one millimeter [2].

The cross-section of the organ of Corti can be divided into two components, a fluid-filled space on the neural side of the organ, and a portion representing the body of Hensen cells on the abneural side (see Fig. 1(B)). The cross-sectional area of each component needs to be conserved separately: the fluid space because of the argument above, and the Hensen cells because their cytoplasm cannot escape longitudinally.

### Description of the geometric model

The motion of the cochlear partition can be decomposed into a passive component, where all structures follow the sound-evoked displacement of the basilar membrane [24], and an active component that involves internal deformation of the organ of Corti caused by outer hair cell forces. Here we seek to determine the motion of various structures of the organ of Corti—in particular of the Hensen cells, the reticular lamina, and the outer hair cells—relative to the basilar membrane.

We use the constraint of a conserved cross-sectional area to estimate the active deformation of the organ of Corti from its geometry. The length change of outer hair cells is characterized by its relative contraction *∊*, such that the length of an outer hair cell is given by *L*_OHC_(*∊*) = (1 — *∊*)*L*_OHC,0_, where *L*_OHC,0_ is the resting length of the cell (Fig. 1(B)). A length change of the outer hair cells can result from electromotility as well as from hair bundle motility that can exert force on the reticular lamina [25–27]. Experiments using electrical stimuli on isolated and unloaded outer hair cells indicate that the magnitude of the outer hair cell contraction is ∣*∊*∣ ≲ 0.02 [4]. Other anatomical elements of the organ of Corti are assumed to have constant length, except for the Deiter’s cells and the contour of the Hensen cells. Motion of the reticular lamina can be approximated as pivoting about the top of the pillar cells, which is why we lump the three rows of outer hair cells and Deiter’s cell in a single, effective row, located near the third row of outer hair cells.

Since we consider small deformations only, we assume linear relationships between the length change of the outer hair cells and the length L_DC_(*∊*) of the Deiter’s cells as well as the length *L*_HC_ (*∊*) of the contour of the Hensen cells. We can therefore write *L*_DC_(*∊*) = (1 + *∊*Δ)*L*_DC_,_0_ and *L*_HC_(*∊*) = (1 + *∊*Γ)*L*_HC_,_0_ with the resting lengths *L*_DC_(*∊* = 0) = *L*_DC_,_0_ and *L*_HC_(*∊* = 0) = *L*_HC,0_. The two parameters Δ and Γ quantify the extent to which the Deiter’s cells and the Hensen cell contour change their length as a result of an outer hair cell length change, respectively. In the following, we will refer to them as extensibilities. Note that we have introduced Δ and Γ as purely geometrical parameters; they do not correspond in a simple way to material properties of the Deiter’s or Hensen cells alone. Rather, they are the result of the complex interplay of the material properties and the geometrical arrangement of all the different elements that comprise the cochlear partition and resist deformation through outer hair cell forces. Their values are therefore *a priori* unknown.

We assume that the Deiter’s cells can pivot around their attachment on the basilar membrane and that they do not bend. The arc of Hensen cells is treated as a thin elastic body that deforms around a preferred shape, characterized by its local curvature along its length. Details of the model calculations are given in the *Materials and Methods*.

### Determining parameter values

We characterize the motion of the deforming organ of Corti by the motion of specific points along the arc of Hensen cells (Fig. 2). Our model shows that, depending on the values of the Deiter’s cell extensibility Δ and the extensibility Γ of the Hensen cell contour, different types of motion can occur. Which of these does occur in experiments?

**Fig 2.**
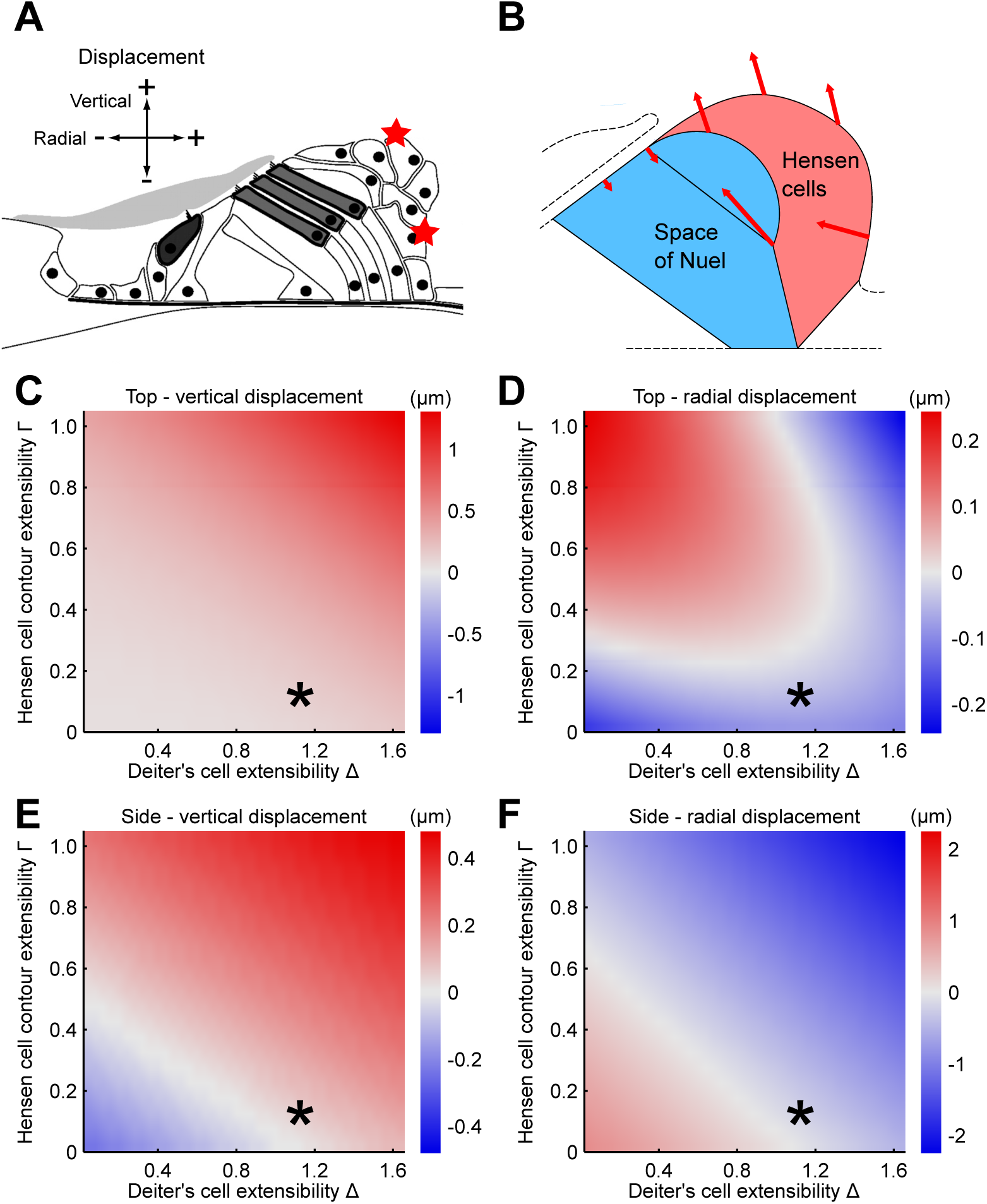
Predicted motion of the Hensen cells for different parameter values. (**A**) We characterize the motion of the Hensen cells through the radial and vertical displacements of two points on the top and on the side of the Hensen-cell contour (red stars). (**B**) The motion pattern predicted by our model through comparison with experimental data involves large displacement of the Hensen cells as well as of the base of the outer hair cells way from the basilar membrane upon outer hair cell contraction. The basilar membrane is assumed to be fixed. (**C**-**F**) Vertical and radial displacement of the two points on the Hensen-cell contour for a hair-cell contraction *∊* = 0.005 and different choices of the mode parameters Δ and Γ. The parameter values that are identified as biologically realistic through comparison with experimental data are indicated through an asterisk and are used in (B). (**C**) The top of the organ consistently moves away from the basilar membrane when the outer hair cells contract. (**E**) The radial displacement of the upper point shows a more complex behaviour: both motion towards and away from the stria vascularis can occur under outer hair cell contraction, depending on the values of the model parameters. (**D**-**F**) The direction of both the vertical and the radial motion of the lateral point depend on the values of the model parameters as well. However, this motion was not experimentally accessible.

### Experimental measurement of the radial motion of the Hensen cells

To determine the parameter values that yield realistic motion patterns in our model, we performed *in vitro* experiments to establish the radial motion of the Hensen cells upon hair cell contraction. Motion of the organ of Corti was induced by applying currents inside scala media. In control experiments where we perfused scala tympani with either sodium salicylate or MET-channel blockers, we have demonstrated previously that this motion is indeed due to prestin-dependent outer hair cell forces [21]. The resulting displacements were measured by confocal imaging of the reticular lamina, as well as interferometry.

**Fig 4.**
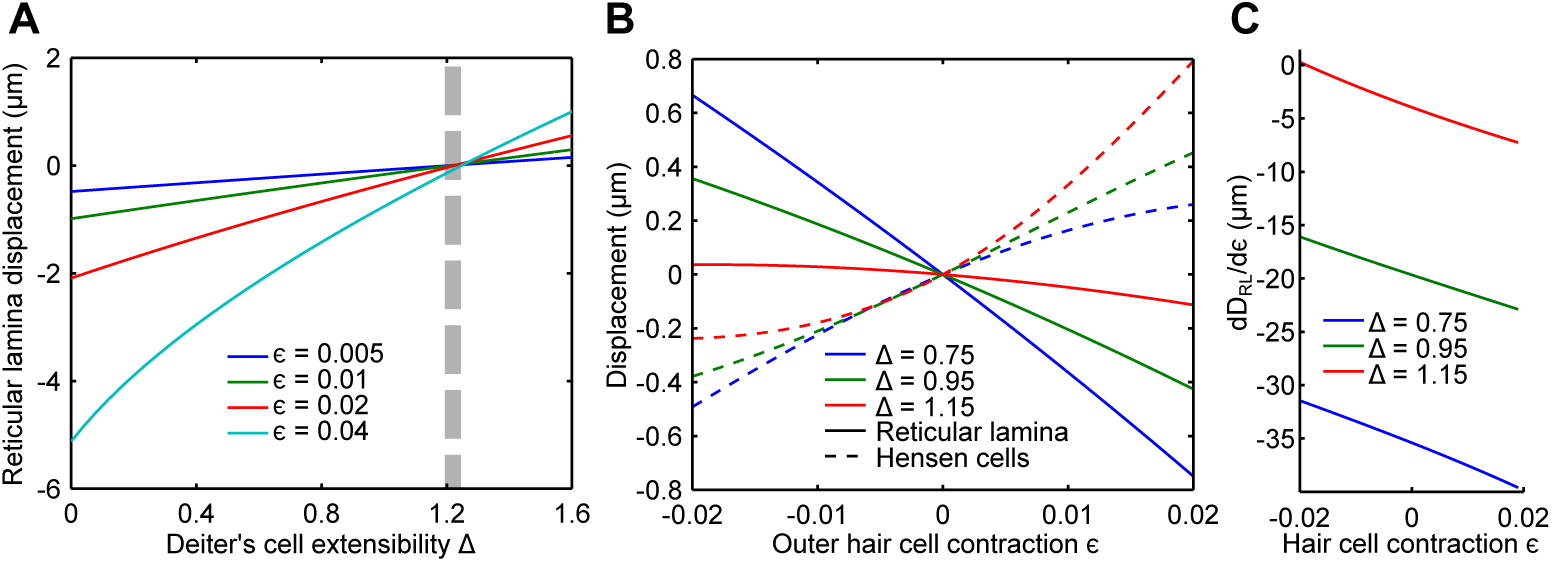
Predicted motion of the reticular lamina for different parameter values. (**A**) A small value of the Deiter’s cell extensibility Δ leads to a large reticular-lamina displacement. At a critical extensibility Δ*_C_* ≈ 1.2 (dashed strip) the displacement vanishes. The critical extensibility Δ*_C_* varies slightly with the outer hair cell contraction *∊*. (**B**) The Deiter’s cell extensibility Δ strongly influences the relation between reticular-lamina displacement (dashed) and Hensen-cell motion (solid) for the model parameter Γ = 0.1 as identified from comparison with experiments. The Hensen-cell motion for the model parameter Δ = 1.15 (red) is in very good qualitative agreement with experimental results of *in vitro* Hensen cell motion under applied current [21,29]. Both the motion of the Hensen cells and of the reticular lamina depends nonlinearly on the contraction ∊ of the outer hair cells, and this nonlinearity is particularly pronounced for a Deiter’s cell extensibility Δ close to the critical value Δ*_C_*. (**C**) The nonlinear dependence in the reticular-lamina motion *D_RL_* on the contraction of the outer hair cells *∊* implies that the absolute value of the derivative of *D_RL_* with respect to the contraction *∊* varies with *∊*. The relative change is particularly strong for a large extensibility Δ of the Deiter’s cells, which has important functional implications.

Experiments using confocal microscopy show that the reticular lamina often exhibits a pivot point between the second and third row of outer hair cells when an external current is applied (Fig. 3(A); data replotted from a previous study [21]). Here, we used 50- or 100-ms long current steps switching rapidly from negative to positive current (as depicted also in Fig. 6(A)) with amplitudes of up to 30 *μ*A. The displacement of the third row of outer hair cells presumably follows the motion of the adjacent Hensen cells and suggests a motion predominantly towards scala vestibuli. Only a very small radial component is found which on average points toward the center of the cochlear spiral (Fig. 3(B,C)).

**Fig 3.**
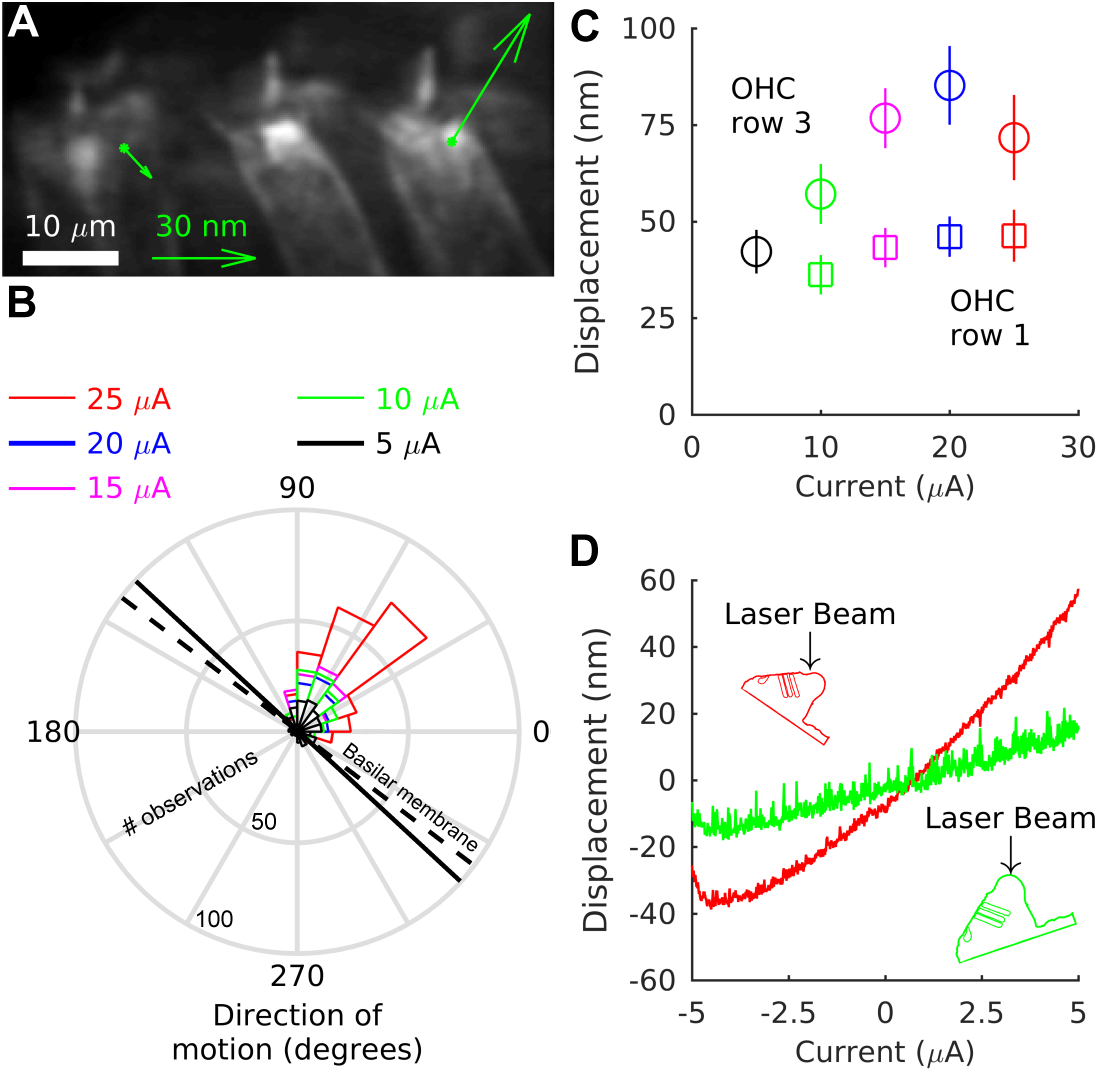
Direction of motion of the Hensen cells. (**A**) Confocal microscopy shows the motion of the reticular lamina when a negative externally-applied current is switched to a positive current of equal magnitude, causing contraction of the outer hair cells. The green arrows show the displacement for the first and third row of outer hair cells (the displacement of the second row was similar to the first row). A pivot point emerges between the second and third row of outer hair cells: the first and second row move towards the basilar membrane whereas the third row moves away from it, following the displacement of the Hensen cells [21]. (**B**) Direction of displacement of the third row of outer hair cells. In this angle histogram, 0° corresponds to motion directed to the right in the image shown in panel A. According to morphometric measurements by Kelly, the basilar membrane is inclined by 37.26° on average with respect to the reticular lamina (dashed line) [28]. Our own measurements from anatomical 3D-reconstructions indicate that this inclination is slightly, but significantly, larger in the undamaged organ of Corti of our in *vitro* cochlear preparation (42.77° ± 6.43°, continuous black line; *N* = 13, *p* = 0.009 by two-tailed t-test, *t* = 3.09, d.f.= 12.). (**C**) The first row of outer hair cells (squares) moves only little. The larger displacement of third-row outer hair cells (circles) mirrors the large displacement of the Hensen cells. Error bars indicate the standard error of the mean from the different measurements. Data in (A-C) are from 683 measurements from 15 preparations for the first row of outer hair cells, and from 905 measurements from 18 preparations for the third row of outer hair cells. (**D**) The radial component of the Hensen-cell displacements was measured directly by tilting the preparation with respect to the interferometer beam. Representative data from one preparation show that the largest motion occurs in a direction with a small component towards the modiolus (red) for positive current injections, consistent with the reticular-lamina data shown in (A,B). Consistent results were obtained from four additional preparations.

We then used interferometry to determine the radial motion further away from the reticular lamina. While the side of the organ of Corti facing the stria vascularis, a dense capillary network along the outer wall of the cochlea, was not accessible, it was possible to estimate radial motion at the top of the organ by tilting the preparation with respect to the interferometer beam (Fig. 3(D)). These measurements suggest that the Hensen cells move with a small component towards the modiolus and away from the stria vascularis when positive current is injected, while the major displacement component is directed towards scala vestibuli, consistent with data from the reticular lamina.

### Displacements of reticular lamina and Hensen cells as functions of outer hair cell contraction

The direction of reticular-lamina displacement that is consistent with the conservation of the cross-sectional area of the organ of Corti depends on the value of the Deiter’s cell extensibility Δ. For values smaller than a critical value Δ*_C_* ≈ 1.2, the reticular lamina is pulled towards the basilar membrane upon outer hair cell contraction, and pushed away from it for values larger than Δ*_C_* (Fig. 4(A)). The latter is inconsistent with experimental observations [17–22]. Thus, the value of Δ must be less than the critical value Δ*_C_* ≈ 1.2. The model further predicts that for values of Δ that are smaller but near the critical value Δ*_C_*, both the displacements of the reticular lamina and the Hensen cells depend nonlinearly on the outer hair cell contraction *∊* (Fig. 4(B,C)). The displacements plateau for outer hair cell elongation (negative values of *∊*), but continue to grow upon contraction (positive values of *∊*). This is comparable to the experimentally observed dependence of Hensen-cell displacement on an externally-applied current [17,21]. Moreover, the ratio between the amplitude of displacement of the Hensen cells and of the reticular lamina resembles that found experimentally, where the larger motion occurs at the Hensen cells [21] (see also Fig. 3(C)). We conclude that the Deiter’s cell extensibility is about Δ = 1.15.

The polarity of the radial Hensen-cell motion corresponds to model parameter values for which radial displacement at the top of the organ upon contraction of outer hair cells is negative. Because of the constraint Δ < 1.2 this restricts the Hensen cell contour extensibility to Γ ≲ 0.2 (Fig. 2(D)). A small value for Γ is plausible, but too small values violate our assumption of conserved cross-sectional area of the organ of Corti for physiological values of outer hair cell contraction. For these small values, no solution to the model equations exist when the hair cell contraction ∣*∊*∣ ≤ 0.02. The biologically-realistic value of the extensibility of the Hensen cell contour is thus Γ = 0.1.

Hence, we find that the parameter values Δ = 1.15 and Γ = 0.1 yield excellent agreement with qualitative features of experimental data, whereas all other regions in the parameter space yield qualitatively different behaviour that does not match the observations. We use these parameter values for our further analysis.

### Displacement of outer hair cells

Having constrained both free parameters of our model, we compared the resulting model predictions to additional known features of apical micromechanics. Recent *in vitro* experiments have shown that outer hair cells essentially pivot around their attachment at the reticular lamina when stimulated electrically [21]. Outer hair cells were first subjected to a negative current, yielding a reference state, and then to a positive current of equal magnitude. The change in current leads to contraction of the outer hair cells which were found to rotate the cell’s base towards the stria vascularis (Fig. 5(A)). The angle of this rotation was quantified for different amplitudes of electrical stimulation and was found to increase linearly for small stimulation amplitudes but to saturate at larger ones [21] (Fig. 5(B)).

**Fig 5.**
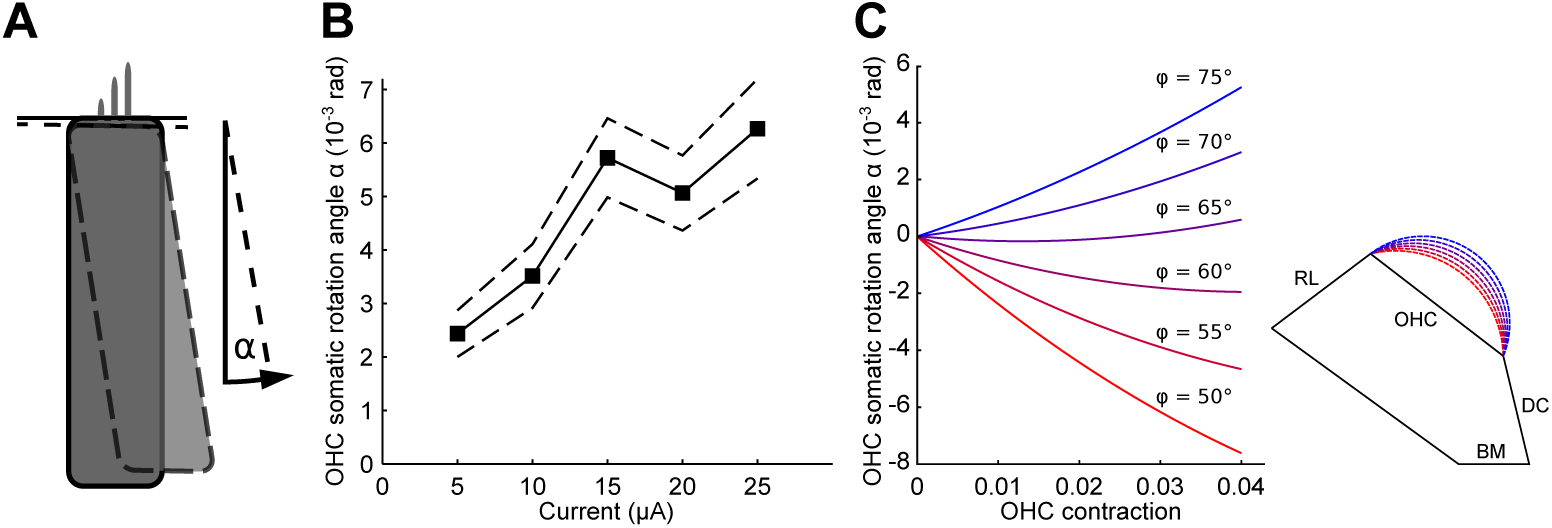
The effect of geometry on outer hair cell displacement. (**A**) *In vitro* experiments show that, under current stimulation, the outer hair cell pivots around its apex. This motion can be characterized by the angle *α* of somatic rotation. (**B**) The somatic rotation angle *α* increases with the size of the current stimulation and saturates for high values (black line, data from 1639 measurements from 24 preparations; dashed lines indicate the 95% confidence intervals). The displacement is directed towards the stria. The data are reused from earlier experiments [21]. (**C**) The predicted angle *α* of rotation of the outer hair cells varies with the size of the arc of the outer tunnel. A positive angle corresponds to a counter-clockwise rotation. The largest arc of the outer tunnel (blue) represents the biologically-realistic geometry and implies an outer hair cell rotation that agrees well with measurements in both direction and magnitude.

Our model shows that the reticular lamina moves much less than the length change of the outer hair cells which is consistent with the essentially rotational motion of the outer hair cells found experimentally. Direction and amount of the rotation depend on the size of the outer tunnel of Corti as parametrized by the angle *φ* between the outer hair cells and the arc of the outer tunnel (Fig. 1(B), Fig. 5(C)). For simplicity, we here consider the reticular lamina as fixed and regard the organ at the hyporpolarized state of the outer hair cell as the reference position. The amount of contraction considered in Fig. 5(C) therefore ranges from *∊* = 0 to approximately *∊* = 0.04, rather than from *∊* = −0.02 to *∊* = 0.02 as before. Our model correctly predicts the direction of outer hair cell rotation when the outer tunnel is large, which agrees with the geometry commonly seen in micrographs: the realistic geometry is arguably the one where the outer tunnel arc lies in almost tangential continuation of the reticular lamina. Furthermore, the amplitude of the applied current can be related to the amplitude of the length change of the outer hair cells if we assume that the saturation observed for high currents corresponds to the maximal contraction of the outer hair cells of about 4% [4, 20]. The predicted rotation angles for a realistic geometry (the light blue curve in Fig. 5(C)) are then in good quantitative agreement with the experimental data (Fig. 5(B)).

### Displacement of Hensen cells at different focal levels

While our model does not explicitly describe displacements of internal points of the Hensen cells, it suggests a motion pattern in which the entire body of Hensen cells is essentially displaced as one by the contracting outer hair cells with little internal deformation. As a consequence, structures at different depths within the organ are expected to show approximately constant vertical displacement, and the displacement decreases only close to the basilar membrane. In particular, the direction of displacement remains the same throughout the entire height of the organ. In contrast, if the cross-section of the organ of Corti were to change, such as through fluid being pressed into the outer tunnel, vertical displacement would vary markedly and change direction as a function of depth.

We interferometrically determined current-evoked displacements from positions at different depths. We found that the direction of Hensen-cell displacement, as well as the displacement amplitude, vary little with depth (Fig. 6(A)). While the direction of the displacements with respect to the applied current was consistent in all preparations, the amplitude of the evoked displacements varied considerably between preparations, as well as with time in a given preparation. For this reason, results shown in Fig. 6(B) have been normalized to the average displacement at the surface of the Hensen cells. The displacement amplitude exhibited a small but significant decrease with increasing depth (13% on average; a linear mixed model reveals a negative slope of −0.0008/*μ*m in normalized displacement units, *p* = 0.0014, *t* = —3.22, d.f.= 532; data from 540 measurements from 7 preparations). This agrees with our model that revealed that the counter-intuitive direction of Hensen-cell motion under electrical stimulation is due to large motion at the bases of outer hair cells.

**Fig 6.**
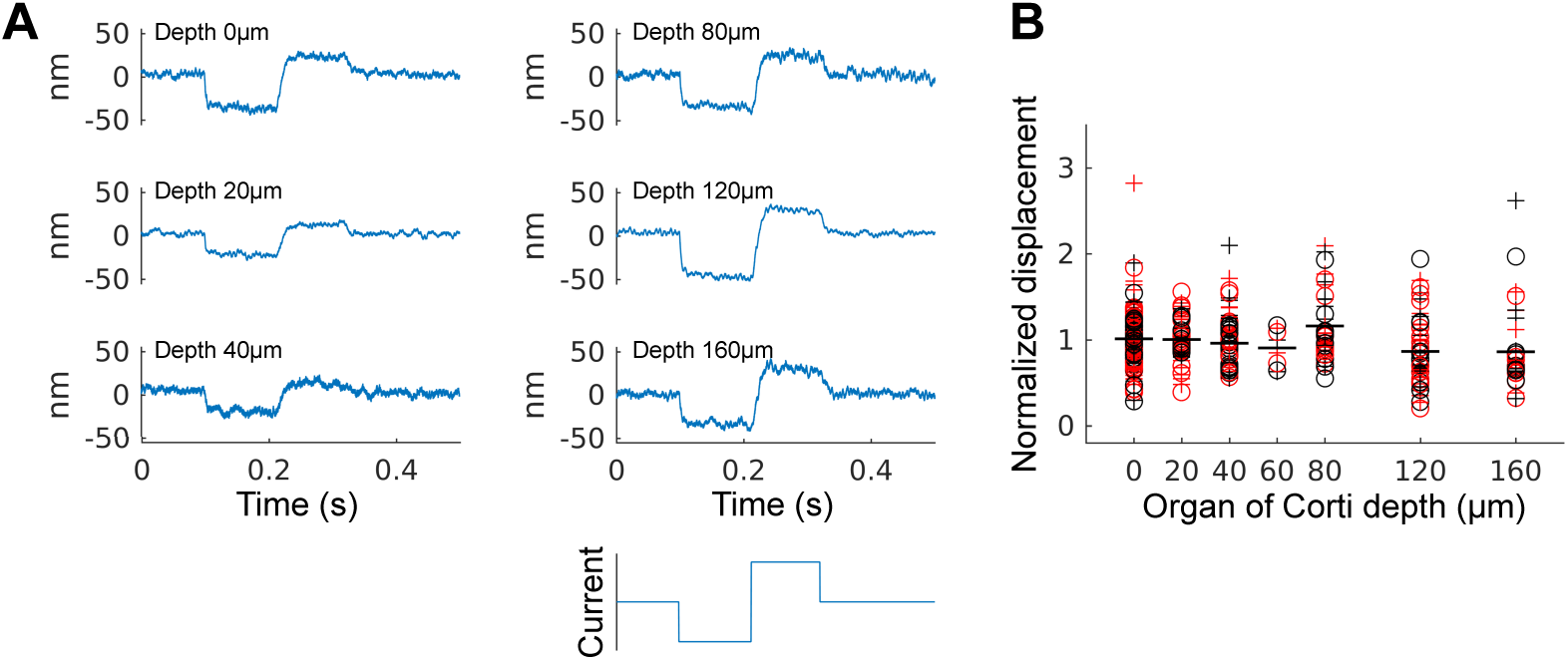
Displacement at different depths in the organ of Corti under current stimulation. (**A**) Representative recordings at different depths under the arc of Hensen cells under negative and positive current stimulation (bottom). (**B**) Displacements of cumulative data from various pulse protocols. The data have been normalized with respect to the average displacement at the surface of the organ, due to high variability in absolute values between preparations, and with time for a given preparation. The displacement vary only little with increasing depth of measurement, neither for positive (’(+)’) nor negative (‘(o)’) currents, and neither in the presence (red) or absence (black) of sound stimuli. Mean values for each set of data with respect to depth only are shown by a flat line.

As the measurement became increasingly noisy with increasing depth inside the tissue, we were not able to determine the location of the basilar membrane. The fact that large displacement amplitudes persist with depth suggests, however, that some basilar membrane motion occurs underneath the Hensen cells. In contrast, such motion was not detectable in the portion of the basilar membrane lateral to the organ of Corti. This is consistent with recent in *vivo* measurements obtained using optical coherence tomography [17].

### Functional implications of the predicted reticular-lamina motion

Inner hair cells are responsible for detecting the mechanical sound vibrations and transducing them into electrical signals that are then forwarded to the brain. The hair bundles of the inner hair cells are deflected by oscillatory fluid flow between the reticular lamina and the tectorial membrane, whose magnitude is dependent on the vibration amplitude of the reticular lamina, at least for frequencies up to 3 kHz [18]. Therefore, the nonlinear reticular-lamina displacement upon length change of the outer hair cells that is predicted by our model has striking consequences for inner hair cell stimulation (Figs. 4(B,C)).

Sound vibration at a frequency *f* leads to an oscillating length change of the outer hair cells around some resting position *∊*^(0)^:

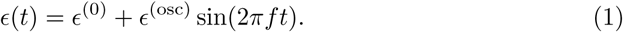

This length change elicits an oscillating reticular-lamina motion *D*_RL_ (*t*) at an amplitude 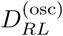 around the steady displacement 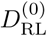 that is set by the outer hair cell’s steady contraction *∊*_0_:

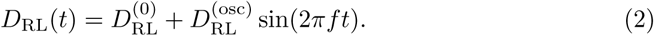

**Fig 7.**
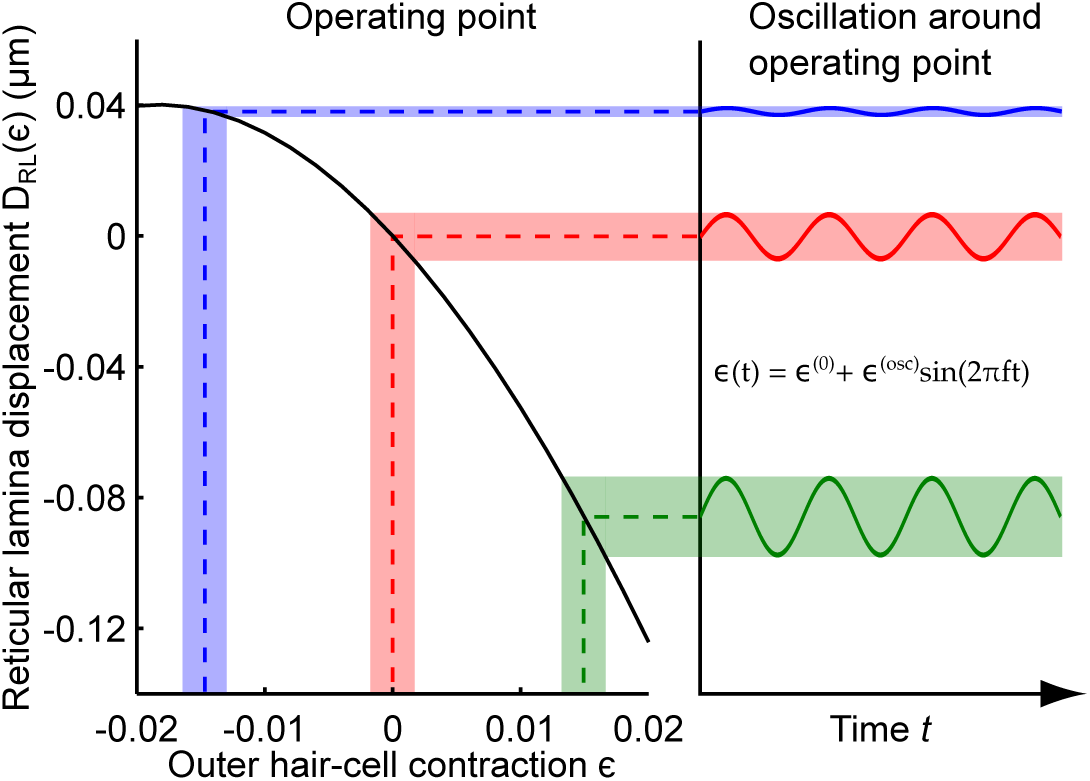
Resting length of outer hair cells can modulate reticular-lamina vibration. How much reticular-lamina vibration is evoked by an oscillatory length change of the outer hair cells depends critically on the operating point set by the static length change of the outer hair cell. An oscillatory length change of an outer hair cell around an elongated state, characterized by a negative value of *∊*, leads to only a very small motion of the reticular lamina (blue). The vibration of the reticular lamina becomes increasingly larger for outer hair cells that oscillate around a progressively more contracted length (red and green).

The amplitude of an oscillating length change of an outer hair cell for sound pressures in the hearing range is small [30], ∣*∊*^(osc)^∣ ≪ 0.02. The amplitude of the resulting reticular-lamina vibration 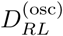 can thus be approximated by a linear expansion around the resting amplitude 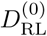:

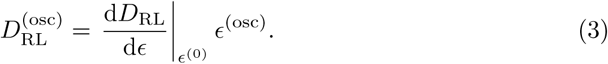

For the Deiter’s cell extensibility Δ = 1.15 that we identified above, the derivative of reticular-lamina displacement with respect to hair-cell contraction, dD_RL_/d*∊*, varies monotonically from approximately zero at a resting length change *∊*^(0)^ = −0.02 of the outer hair cells to a value of approximately −7 at *∊*^(0)^ = −0.02. As a result, an oscillating length change of the outer hair cells around a maximally elongated resting length defined by *∊*^(0)^ = −0.02 produces virtually no oscillation of the reticular lamina. On the other hand, a vibration of the outer hair cell length around a maximally contracted resting length defined by *∊*^(0)^ = −0.02 leads to a seven-fold larger vibration of the reticular lamina (Fig. 7). The resting length of the outer hair cells can thus sensitively determine how much vibration of the reticular lamina is elicited by an oscillating length change of the outer hair cells at low frequencies.

## Discussion

We have developed a model for the deformation of the organ of Corti that is based on the organ’s geometry as well as on the plausible assumption that the organ of Corti near the cochlear apex is incompressible. The model involves only two parameters that are not derived from the geometry, namely the extensibility of the Deiter’s cells and of the outer edge of the Hensen cells. Qualitative comparison of model predictions with experimental data highly constrains these parameters, and the resulting model predictions agree excellently with further data on the displacement of the outer hair cells and the vertical vibration at different depths in the organ of Corti. Our argument holds for frequencies at which the organ of Cort’s impedance is dominated by stiffness, rather than viscosity, which holds for frequencies up to several hundred Hertz [31]. This corresponds to the characteristic frequencies found at the apical locations studied here, which is why our results are relevant also for the acoustic response.

Our model generically produces the non-intuitive counterphasic motion of the reticular lamina and the Hensen cells that was recently observed experimentally [18,19,21,22]. Importantly, our analysis suggests that this behaviour does not result from perilymph being pressed against the Hensen cells, as hypothesized recently [22]. Instead, our model and our measurements evidence that the entire body of Hensen cells is being pulled upwards by the contracting outer hair cells. Generally, the experimental data are reproduced if the base of the outer hair cell is allowed to move somewhat more than its apex, such that the largest displacements then occur inside the organ of Corti. Intriguingly, this is corroborated by our own recent in vivo measurements using optical coherence tomography [17].

What is the origin of this internal motion? In our model, the vibrational pattern is achieved through Deiter’s cells that are fairly compliant, at least in response to quasi-static or low-frequency forcing by outer hair cells [32,33]. Alternatively, or in addition, large displacements at the bases of outer hair cells could also occur as a consequence of a locally very compliant basilar membrane [17]. We have not detected basilar-membrane motion lateral of the organ of Corti in response to current stimulation [17,29]. However, our interferometric measurements from different depths inside the Hensen cells indicate that some basilar-membrane motion may be present in a limited region underneath the organ, while the decrease in amplitude with depth suggests that some stretching occurs as well. Conservation of the cross-sectional area of the organ of Corti may then require counterphasic displacement of the arcuate zone of the basilar membrane, as observed by Nuttall et al. in more basal regions in response to electrical stimulation [34]. We did not include this mode of deformation in our model, as no corresponding data are available for the cochlear apex.

Current theories of cochlear function suggest that the mechanical activity of outer hair cells serves to amplify the motion of the basilar membrane [2] or the reticular lamina [35, 36] in order to render faint sounds more easily detectable for the stereocilia of inner hair cells. In this light, it seems surprising that the largest motion would occur in the interior of the organ. However, our geometrical analysis and experiments suggest that this motion pattern is associated with a nonlinear dependence of the reticular-lamina motion on the resting length of the outer hair cells. In consequence, we find that the resting length of the outer hair cells can control the magnitude of vibration of the reticular lamina that is evoked by an oscillating length change of the outer hair cells. Experimental evidence for this effect comes from the observed nonlinear dependence of sound-evoked motion on an imposed endocochlear potential in *vitro* [21]. It has been suggested previously that static length changes of the outer hair cells might influence the operating point of hair bundles [37], or of the micromechanics of the organ of Corti as a whole [38], but the details of such a mechanism have remained unclear. Our analysis shows that the incompressibility of the organ of Corti together with a high level of compliance at the base of outer hair cells yields a novel and intriguingly simple mechanism for the outer hair cells to regulate hearing sensitivity through their static length change. While we have thus shown the availability of such a mechanism, further experimental work and improved imaging techniques are needed to verify it in the living cochlea.

Our geometrical model quantifies the internal motion of the organ of Corti. The actual sound-evoked and active motion of the cochlear partition is a linear combination of the internal deformation and an overall net displacement. While internal motion is due to active amplification by outer hair cells, the net displacement of the organ can be caused both by sound stimulation as well as by the mechanical activity of outer hair cells. In a recently proposed ratchet mechanism, or unidirectional amplification, the outer hair cells may cause only internal deformation of the organ of Corti without displacement of the basilar membrane [25], in agreement with some recent experimental observations [17]. Further modeling that integrates the geometric model presented here with an analysis of the different forces produced by outer hair cells and their effects on the overall motion of the organ of Corti, as well as further experimental results on the linear or nonlinear response of the reticular lamina and the basilar membrane to varying sound intensity, are needed to clarify these issues.

Our findings are particularly relevant for two lines of further research. First, our results could shed new light on the role of the static and frequency-dependent motile response of outer hair cells to acoustic stimulation whose biophysical origin and function in the cochlea remain poorly understood [13,14]. Because our model shows that a sustained length change of outer hair cells can sensitively regulate the reticular lamina’s vibration, the tuned sound-evoked static length changes of outer hair cells can serve as an effective tuning mechanism that can circumvent the poor mechanical tuning of the basilar membrane in the cochlear apex [2]. As set out above, elongated outer hair cells will transfer only little of their oscillating length change to the reticular lamina. The mechanical sound signal elicited by a pure tone may, however, cause outer hair cells at the characteristic position to contract such that their additional oscillatory response to sound is leveraged into a large vibration of the reticular lamina and thus of the hair bundles of the inner hair cells. This effect can thus endow the motion of the reticular lamina with a frequency selectivity that is independent of the mechanical tuning of the basilar membrane which is comparatively poor in the cochlear apex [2].

Second, the discussed principle could present a potential mechanism for efferent medial olivocochlear (MOC) nerve fibers that innervate the outer hair cells to modulate the auditory stimulus [39]. This efferent feedback is thought to play an important role, for instance, in our ability to understand speech in noisy environments. Our results show that efferently-mediated static length changes of the outer hair cells can modify the transfer of outer hair cell activity to reticular-lamina motion. Experiments have indeed observed efferently-induced modifications in the auditory nerve signal that is not found in the mechanics of the basilar membrane, suggesting that inner hair cell stimulation is in part directly due to outer hair cell activity [40]. This effect was present throughout the cochlea, and was particularly prominent in low-frequency regions. A mechanism as the one described here could underlie these observations.

## Materials and Methods

We used detailed morphometric data on the guinea pig’s organ of Corti in the cochlear apex [28,41,42] in conjunction with high-quality micrographs [15] as a basis for the geometrical model (Fig. 1). Relative sizes and orientations of different structures in the organ of Corti show a high level of consistency between the different data sources. The contour of the Hensen cells is represented by a polynomial curve approximating the shape seen in micrographs. Since we assume the reticular lamina to pivot as a stiff rod around its attachment near the inner hair cell [8,9], we have for simplicity lumped the three rows of outer hair cells and Deiter’s cells into a single one, located at the position of the outermost row.

### Model Equations: Deformation of the fluid space

The fluid space of the organ of Corti can be further decomposed into three subcomponents (Fig. 1(B)): the triangle formed by the basilar membrane and the pillar cells (the tunnel of Corti) with cross-sectional area *A*_TC_, a polygonal space defined by basilar membrane, reticular lamina, the outer pillar, the outermost outer hair cell, and the Deiter’s cell (the space of Nuel) with cross-sectional area *A*_SN_, and the outer tunnel adjacent to the Hensen cells with cross-sectional area *A*_OT_. The equations determining the configurational change of the fluid space upon outer hair cell contraction, such that the total cross-sectional area of the organ of Corti remains conserved, as as follows.

The elements shaping the tunnel of Corti are comparatively stiff [43], so that we assume its shape to be unaffected by outer hair cell forces and accordingly *A*_TC_ = const..

The outer tunnel is assumed to be of circular shape with an arc length a. When it deforms upon outer hair cell contraction, it deforms into another circular segment with the same arclength. This is motivated by the fact that its outer wall is supported by stiff polymer cables [16]. Let *φ* be the angle between the outer-tunnel arc and the adjacent outer hair cell (Fig. 1(B)). The length *L*_OHC_ of the outer hair cell, corresponding to the chord length of the circular segment representing the outer tunnel, is given by

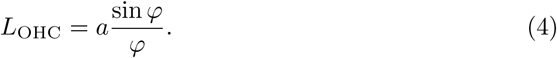

The area *A*_OT_ of the outer tunnel as a function of *a* and *φ* is

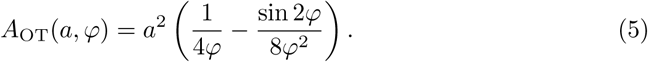

Let now *L*_OHC,0_ be the initial length of the adjacent outer hair cell and assume it changes to *L*_OHC_(*∊*) = (1 − *∊*)L_OHC 0_ for some small value *∊* with ∣*∊*∣ ≪ 1. We refer to this variable as the *contraction* since *∊* > 0 corresponds to a shortening and *∊* < 0 to an elongation of the outer hair cell. If initially *φ*(0) = *φ*_0_, then we find the new angle *φ*(*∊*) using equation (4), which yields the relation

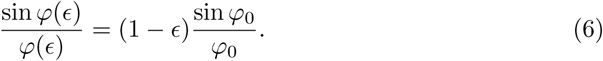

To account for the possibility that the Deiter’s cell might deform elastically [33] rather than just rotate in response to outer hair cell contractions, we introduce the *Deiter’s-cell extensibility* Δ as a free parameter such that the length of the Deiter’s cell is given by *L*_DC_(*∊*) = (1 + *∊*Δ)*L*_DC_,_0_. A linear relation is justified since experiments on isolated outer hair cells have established that the outer hair cell contraction is small, ∣*∊*∣ ≲ 0.02 [4].

In contrast, we assume that the reticular lamina is stiff enough such that it neither stretches nor bends. Its motion is thus constrained to rotation around its attachment at the apex of the outer pillar cell.

The area *A*_SN_ of the polygonal space of Nuel is found using the general formula for the area of a polygon with *N* vertices *x_i_* = (*x_i_*,*y_i_*),

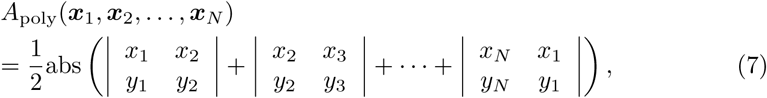

where abs(·) denotes the absolute value and ∣·∣ are determinants. In our model, the only vertices that are not fixed by assumption are the two endpoints ***a*** (the cellular apex) and ***b*** (the base) of the outer hair cell, so that *A*_SN_ = *A*_SN_(***a***, ***b***). We see that given an outer hair cell contraction *∊*, we require four equations to determine the deformed configuration of the fluid space. Three equations follow from the conserved lengths of the reticular lamina, the outer hair cell, and the Deiter’s cell. The fourth equation results from the constant-area condition and reads

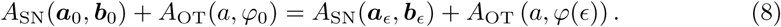

The system of equations is then solved for the unknown endpoints of the outer hair cell ***a***_*∊*_ and ***b***_*∊*_.

Note that in all our calculations we consider only internal deformation of the organ of Corti and keep the basilar membrane fixed as a reference. A basilar-membrane vibration such as elicited by sound can be superimposed to the motion considered here.

### Shape deformation of the Hensen-cell body

The Hensen cells form the abneural portion of the organ of Corti that runs approximately along the midline of the basilar membrane. Rather than modelling the detailed mechanics of the Hensen cells, we take a simplified approach and model the contour of the Hensen cells as a thin elastic body deforming around a preferred shape. The allowed deformations are constrained by the requirement that the cross-sectional area of the Hensen-cell region remains constant. Hence, the deformations are determined by minimizing with appropriate boundary conditions a functional of the form

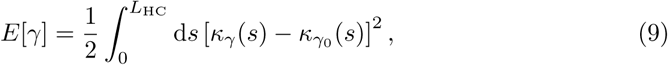

where *L*_HC_ is the length of the contour, *γ*(*s*) and *γ*_0_(*s*) are arbitrary arc-length parametrisations of the deformed and initial contours, respectively, and *κ_γ_*(*s*) denotes the signed curvature given by

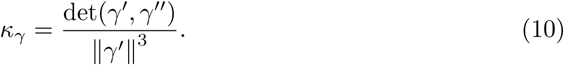

The expression (9) is in fact a one-dimensional version of the Helfrich Hamiltonian used to determine the shape of elastic membranes with spontaneous curvature [44]. This way, however, the contour is implicitly assumed to be inextensible. Clearly, since the contour is the outer boundary of a viscoelastic body, this need not be true and its length may change from an initial length *L*_HC_,_0_ to a new length *L*_HC_(*∊*) = (1 + *∊*Γ)*L*_HC,0_. The amount of length change is determined by the mechanical properties, as well as the geometry of the Hensen cells and is not *a priori* known. The extensibility Γ thus presents another free parameter of our model. To take the length change of the contour into account, we modify the above functional to

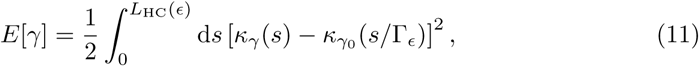

where we used the shorthand notation Γ_*∊*_ = 1 + *∊*Γ. We hereby make the simplifying assumption that the contour is stretched uniformly along its length. As Γ_*∊*_ is always close to one, we also make the approximation of comparing curvatures between the position *s* of the new curve and *s*/Γ_*∊*_; of the initial curve.

It is convenient to parametrize the contour in terms of its local angle *ϕ*(*s*) with respect to an arbitrary but fixed axis, which we take to indicate the *x*-axis in a cartesian coordinate system (Fig. 1(B)). In this coordinate system, the curve is given as *γ*(*s*) = (*x*(*s*),*y*(*s*)) with

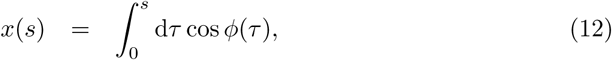

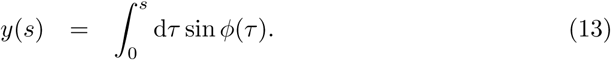

The curvature now takes the very simple form

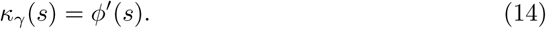

The endpoint *s* = 0 corresponds to the fixed outer edge of the Hensen cell body and lies at the origin. The position (*x*_e_, *y*_e_) of the endpoint at *s* = *L*_HC_(*∊*), coinciding with the apex of the outer hair cell, is determined from the deformation of the fluid space of the organ. This introduces two additional constraints to the variational problem. Extending accordingly equation (11), our complete functional reads

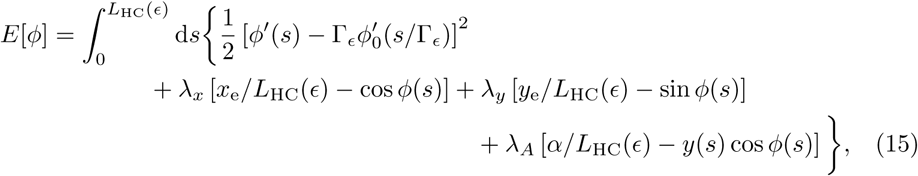

where λ*_x_*, λ*_y_*, and λ*_A_* are Lagrange multipliers to account for the endpoint and area constraints, and *α* is an appropriate constant to ensure the constant-area condition. We require two boundary conditions for the angular dependence *ϕ*(*s*). On the abneural side, *i. e.* at *s* = 0, there is no evident physical constraint imposed on *ϕ*. The appropriate boundary condition is then given by the natural boundary condition 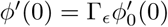. At *s* = *L*_HC_(*∊*), where the Hensen cells join the reticular lamina, we require that the angle between the contour and the outer tunnel arc remains constant. For an appropriate value *ϕ*_e_ the boundary condition hence reads *ϕ*(*L*_HC_(*∊*)) = *ϕ*_e_.

To solve the problem computationally, we derive it in discretized form. For convenience, we divide the arc length *L*_HC_(*∊*) of the contour into *N* equal elements of finite size Δ*s*. The function *ϕ*(*s*) becomes an (*N* + 1)-dimensional vector *ϕ* = (*ϕ_i_* = *ϕ*(*s_i_*)) of function values at the discrete locations *s* = (*s_i_*) along the contour. Introducing **λ** as the vector of Lagrange multipliers and *ϕ_0_* = (*ϕ*_0_*_i_* =*ϕ*_0_(*s_i_*/Γ*_∊_*)) as the vector of initial function values, the functional (15) above becomes the Lagrange function

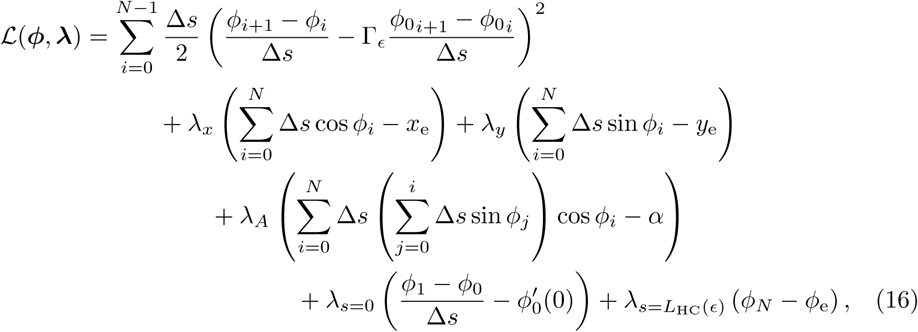

where the boundary conditions for *ϕ*(*s*) at *s* = 0 and *s* = *L*_HC_(*∊*) appear as constraints with corresponding Lagrange multipliers λ_*s*=0_ and λ_*s*_=*L*_HC_(_*∊*_). The optimizing vector *ϕ*_opt_ is finally found as the solution to the system of nonlinear equations

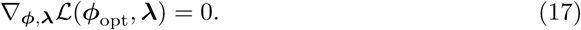

Together with the equations derived for the deformation of the fluid space, this is readily solved iteratively employing the MATLAB routine `fsolve` and using the initial configuration as initial guess for the solution.

## Experimental Methods

Young pigmented and albino guinea pigs of both sexes weighing 200 to 400 g were used in the current study. The animals were housed at six animals per cage and a 12-hour light/dark cycle. Using procedures approved by the local ethics committee (permit N32/13), the temporal bones were removed, attached to a custom holder, and the bulla opened to expose the cochlea. The preparation was then immersed in oxygenated tissue culture medium (Minimum Essential Medium, Invitrogen, Carlsbad, CA, USA) and a small opening created over scala vestibuli in the apical turn. This opening provided optical access to the organ of Corti and also allowed the tip of a beveled glass microelectrode to be pushed through the otherwise intact Reissner’s membrane. The electrode was used throughout the experiment to monitor the sound-evoked potentials produced by the sensory cells. Data collection was aborted if these potentials underwent sudden changes, or if their initial amplitude was abnormally low. The electrode was also used for injecting electrical currents into scala media. The currents were generated by an optically isolated constant current stimulator (A395, World Precision Instruments, Sarasota, FL, USA). For our different experiments, we used either linear current ramps or current steps as stimuli, with durations between 50 ms and 100 ms and amplitudes of up to 30 *μ*A. Scala tympani was continuously perfused with oxygenated tissue culture medium at a rate of ~ 0.6 ml/h, starting within 10 minutes of decapitation, and the perfusion system was also used to introduce the dye RH795 (5 micromolars, Biotium, Howard, CA, USA), which provides fluorescent labeling of the cell membranes of sensory cells and neurons. All experiments were performed at room temperature (21 – 24°C).

### Interferometry and confocal imaging

A displacement-sensitive interferometer (noise floor < 0.1 nm/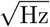 at frequencies above 10 Hz) was used for measuring organ of Corti motion. Cells in the organ of Corti had sufficient optical reflectivity to allow measurements in the absence of artificial reflectors. All displacement data were averaged 10 to 20 times, but traces where the carrier signal of the interferometer had low amplitude were automatically rejected by the Labview-based data acquisition software. The interferometer provides a high-precision measurement of organ of Corti motion, usually from the Hensen cells, but the system can only detect motion directed along the optical axis. To assess the direction of motion, it is necessary to withdraw the electrode, reposition the preparation and the stimulus electrode, and then repeat the interferometric measurements. The complexity of this experiment meant that it was only occasionally successful.

To provide additional data on the direction of electrically evoked motion at the reticular lamina, we used the rapid confocal imaging method described previously [45]. In these measurements, RH795 was applied to the hearing organ as described above, and the dye excited with 488-nm light from a confocal microscope (LSM510, Zeiss, Jena, Germany). A 40x, NA0.8 water immersion lens was used to detect the fluorescence emitted from the dye, using appropriate optical emission filters. To visualize electrically evoked motions, a sequence of 25 - 37 confocal image frames were acquired without interframe delay. The microscope generated a +5V electrical pulse each time a pixel was acquired; this clock signal was used to drive the generation of the electrical stimulus, a square wave at a frequency of 5 Hz. Since the electrical stimulus is applied at a frequency many octaves below the best frequency of the recording location, it is considered static. The method of stimulus generation means that the exact phase of the stimulus with respect to each individual pixel is known, which makes it possible to reconstruct the motion of the sensory cells using custom Matlab scripts. Processing of the image sequence resulted in a new sequence of images, where each frame was specific for one phase of the stimulus. To quantify the motion seen in these image sequences, we used a wavelet-based optical flow calculation method described earlier [46]. Detailed performance evaluations of these techniques [45] have shown that cochlear motion patterns can be accurately quantified down to a motion amplitude of approximately 0.5 pixels. The pixel size was adjusted to allow measurement of motions down to approximately 30 nm.

## Acknowledgments

This research was supported in part by the National Science Foundation under Grant No. NSF PHY11-25915.

